# Variable training but not sleep improves consolidation of motor adaptation

**DOI:** 10.1101/259671

**Authors:** Benjamin Thürer, Frederik D. Weber, Jan Born, Thorsten Stein

## Abstract

How motor memory consolidates still remains elusive. Motor tasks’ consolidation were shown to depend on periods of sleep, whereas pure non-hippocampal dependent tasks, like motor adaptation, might not. Some research suggests that the mode of training might affect the sleep dependency of motor adaptation tasks. Here we investigated whether sleep differentially impacts memory consolidation dependent on the variability during training. Healthy men were trained with their dominant, right hand on a force field adaptation task and re-tested after an 11-h consolidation period either involving overnight sleep (Sleep) or daytime wakefulness (Wake). Retesting also included a transfer to the non-dominant hand. Half of the subjects in each group adapted to different force field magnitudes with low inter-trial variability (Sleep-Blocked; Wake-Blocked), the other half with high variability (Sleep-Random; Wake-Random). EEG was recorded during task execution and overnight polysomnography. Motor adaptation was comparable between Wake and Sleep groups, although performance changes over sleep correlated with sleep spindles nesting in slow wave upstates. Higher training variability improved retest, including transfer learning, and these improvements correlated with higher alpha power in contralateral parietal areas. Enhanced consolidation after training might foster the ability to correct ongoing movements by responsive feedback rather than their pre-execution prediction.

## Introduction

The influence of post-learning sleep on motor memory consolidation has been frequently investigated (1). However, the literature shows an inconsistent picture with studies supporting (e.g. 2-4) and not supporting (e.g. 5,6) sleep dependent motor memory consolidation. Many studies, hence, point to a more complex relationship between specific factors of motor tasks and sleep (7, but also see 8) In a recent qualitative literature review (1), researchers identified that motor benefits or stabilizations due to sleep can be seen in explicit sequence learning tasks, specific variants of implicit sequence learning tasks, and specific visuomotor adaptation tasks, with all of these tasks involving hippocampal function to a certain extent. On the other side, it has been suggested that specific non-hippocampal-mediated tasks, like motor adaptation to dynamic perturbations (e.g. force field adaptation, 9), reflect a motor memory process that is purely time-but not sleep-dependent (5), although those results, to the best of our knowledge, have not been confirmed so far.

Beyond sleep’s dependency on specific task aspects, the effects might also depend on the specific training schedule. Several studies showed that motor training under highly unstable conditions, compared to more stable conditions, enhances posttest and transfer performance, suggesting that depending on the training schedule different memory systems are involved (e.g. 10-12). Furthermore, it has been assumed that specifically benefits after variable training depend on sleep (1). This assumption is derived from a study investigating imaginary training which showed that variable but not constant mental training of a motor task leads to sleep dependent memory improvements (13). Moreover, other studies revealed that hippocampal dependency of a motor task changes with the schedule and the amount of training (14-16).

In this study, we assessed the effects of sleep on memory for a force field adaptation task. Specifically, we were interested whether effects of sleep might express depending on the variability of different force fields used during training. For this purpose, subjects were examined either in more stable or highly unstable training conditions, and retested after periods of sleep or wakefulness with the same arm. Since previous work from our lab showed sleep dependent consolidation effects for contralateral transfer (17), we also examined transfer performance on the contralateral hand. We recorded EEG correlates during training, intervening sleep, and during retest, and also aimed to characterize the role of online feedback mechanisms in mediating improvements during movement execution.

## Results

### Behavioral results

All groups adapted to the dynamic force fields and decreased their motor error (quantified by the enclosed area, EA) during Training (Fig. 2a,b, *F*(1,44) = 143.05, *p* < 0.001, *pEta*^*2*^ = 0.765, for ANOVA with factor time (First Training Trials, Last Training Trials)) independent of Sleep/Wake conditions (*F*(1,44) = 0.06, *p* = 0.801, *pEta*^*2*^ = 0.001) or Blocked/Random training conditions (*F*(1,44) = 1.95, *p* = 0.169, *pEta*^*2*^ = 0.042). The Blocked groups adapted faster during Training than the Random groups (time*training, *F*(1,44) = 4.50, *p* = 0.040, *pEta*^*2*^ = 0.093; time*sleep, *F*(1,44) = 1.29, *p* = 0.262, *pEta*^*2*^ = 0.028, for mixed ANOVA with factors training (Blocked, Random), sleep (Sleep, Wake), and time (First Training Trials, Last Training Trials)). Faster learning for Blocked groups was confirmed by FDR corrected *post-hoc t*-tests on Last Training Trials between Random and Blocked groups (*t*(46) = 3.96, *p* = 0.002, *d* = 1.144).

**Figure 1:**
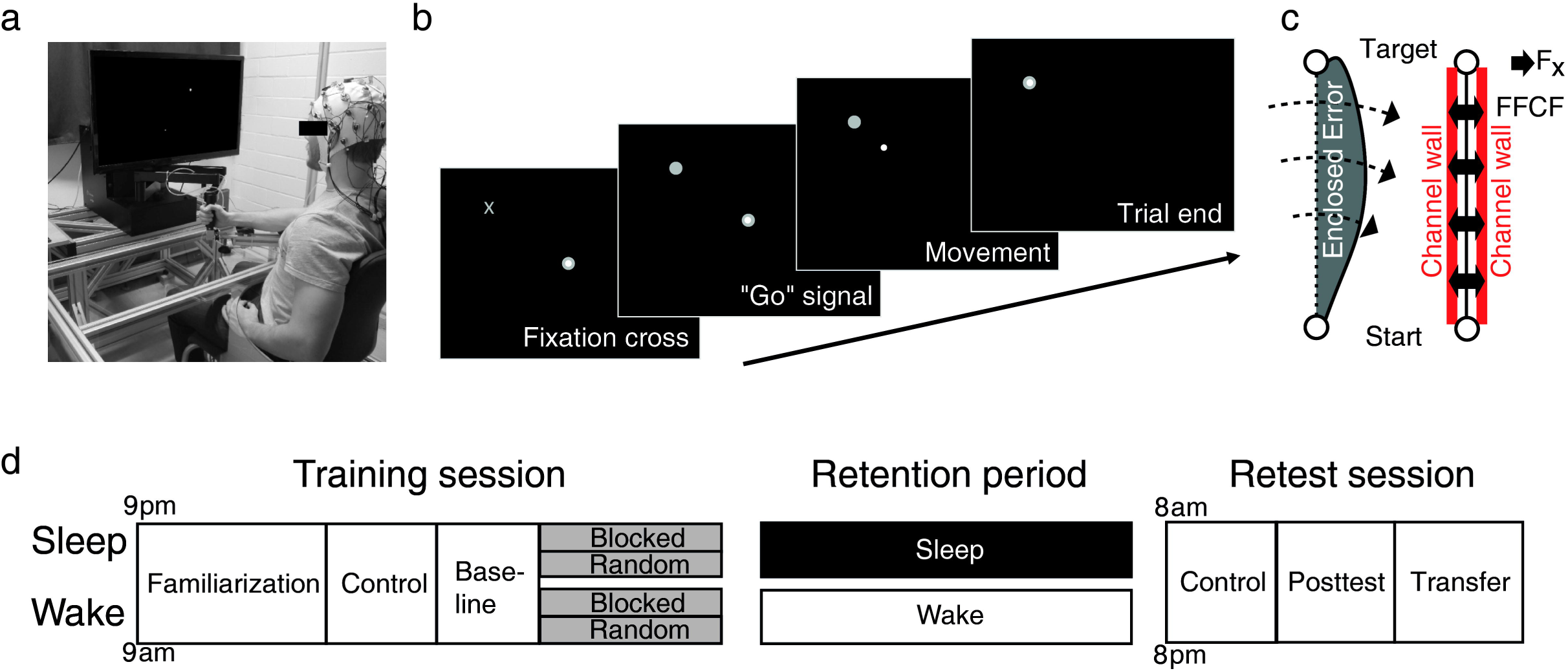
Motor adaptation task and experimental design. (**a**) The motor adaptation task was instrumented by a robotic manipulandum (Kinarm End-Point Lab, BKIN Technologies) with a custom made low friction air-sled system. The robotic manipulandum can induce force fields to perturb participants’ hand movements. During the task, participants’ EEG is recorded. (**b**) Example of one trial from highlighting of the fixation cross to trial termination by reaching the target. (**c**) Sketch of the parameters quantifying the motor error (enclosed area, EA) and motor prediction (force field compensation factor, FFCF). Enclosed area (left) is defined by the area between the trajectory and the direct line between start and target. Arrows indicate the force field direction. The FFCF (right) is computed using the subject’s forces (Fx) directed against virtual channel walls and compared to the ideal force profile to cancel out the perturbation. (**d**) All participant groups had a Training session to train the motor adaptation task with their dominant right hand including Familiarization phase, Control tests (subjective sleepiness, mood, and vigilance), a Baseline, and the force field Training (gray blocks). After a Retention period, Retest performance was quantified in a Posttest and an additional Transfer test on the left hand. Groups differed in their training and consolidation period. The Random groups trained motor adaptation under highly unstable conditions and the Blocked groups under more stable training conditions. The Wake groups trained in the morning and were retested in the evening and the Sleep groups were trained in the evening and retested the following morning after a night of sleep.

**Figure 2:**
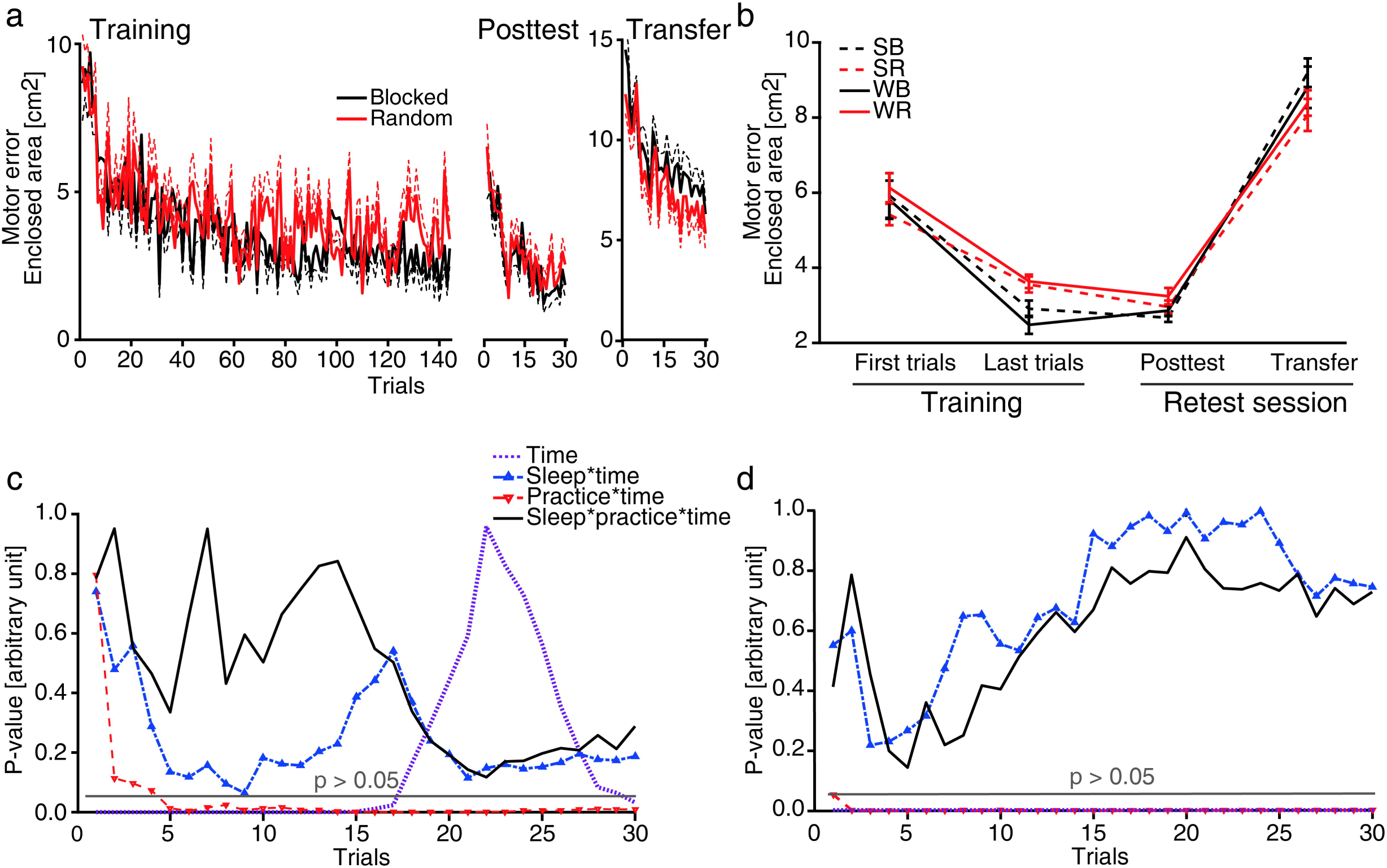
Behavioral results. (**a**) Progress of the mean (± s.e.m) motor error (enclosed area, EA) for the Blocked (black) and Random (red) groups during Training (left), Posttest (middle) and Transfer trials (right). (**b**) Mean (± s.e.m) motor error across 30 trials for each group during First and Last Training Trials, Posttest and Transfer. SB: Sleep-Blocked, SR: Sleep-Random, WB: Wake-Blocked, WR: Wake-Random. (**c**) P-values of different factors and interactions of the mixed ANOVAs investigating a consolidation effect from Training to Posttest and (**d**) Training to Transfer.

Random and Blocked groups started with similar Posttest performance (Fig. 2a,b), showing that the retention from the Last Training Trials (11 hours earlier) to Posttest was worse for the Blocked compared to the Random groups (Fig 2b, retention*training, *F*(1,44) = 6.95, *p* = 0.012, *pEta*^*2*^ = 0.136, for mixed ANOVA with factors retention (Last Training Trials, Posttest) and training (Blocked, Random)), but independent of sleep (retention*sleep, *F*(1,44) = 1.68, *p* = 0.201, *pEta*^*2*^ = 0.037) for any group combination (retention*sleep*training, *F*(1,44) = 1.44, *p* = 0.237, *pEta*^*2*^ = 0.032, for mixed ANOVA with factors retention (Last Training Trials, Posttest), training (Blocked, Random) and sleep (Sleep, Wake)). The benefit of Random over Blocked training manifested quickly during the Posttest from the 5th trial onwards (Fig. 2c). This benefit is also confirmed by Pearson correlation analyses showing that training success (i.e., lower motor error at the end of Training) was inversely related to retention success (percentage of Posttest error related to last Training error, *r* = -0.78, *p* < 0.001, *n* = 48) with this effect weaker in the Random (*r* = -0.51, *p* = 0.01, *n* = 24) than in the Blocked groups (*r* = -0.86, *p* < 0.001, *n* = 24; *p* = 0.018, *z* = 2.36, for the difference using Fisher r-to-z transformation). However, training success in general was moderately predictive and positively correlated with the overall Posttest performance for all groups (*r* = 0.29, *p* = 0.044, *n* = 48; for any group *r* is within 0.167–0.35). This suggests, though initial Training performance might benefit from a blocked training schedule, motor memory retention benefit from a randomized training schedule. These processes were unaffected by sleep.

We further investigated if memory consolidation also enhanced the generalization from the dominant hand (during Training) to the non-dominant (Transfer). All groups performed worse during Transfer testing as compared to the Last Training Trials (*F*(1,44) = 483.56, *p* < 0.001, *pEta*^*2*^ = 0.917). In addition, initial Transfer performance of all groups was worse compared to the initial Training performance (First Training Trials; Fig. 2). This lower initial performance during contralateral transfer learning indicates that participants expected the force field in the opposite direction as during Training (relying on an internal rather than an external representation). This is also supported by the motor prediction (force field compensation factor, FFCF) showing similar force field predictions in the initial Transfer trials for the Blocked and Random groups (Blocked: -15.64 %, SD 16.61 %; Random: -15.62 %, SD 17.85 %, negative sign indicates expectation of opposite force field direction). Motor error in Transfer test was lower in the Random than in the Blocked groups (retention*training, *F*(1,44) = 16.57, *p* < 0.001, *pEta*^*2*^ = 0.274, for mixed ANOVA with factors retention (Last Training Trials, Transfer) and training (Blocked, Random)) and this effect manifested immediately after the first Transfer trial (Fig. 2a,d). Transfer learning effects were independent of sleep (retention*sleep, *F*(1,44) = 0.11, *p* = 0.742, *pEta*^*2*^ = 0.002) and not strongly predictive by training success (for all groups *r* = 0.12, *p* = 0.42, *n* = 48; Random group *r* = 0.14, *p* = 0.54, *n* = 24; Blocked group *r* = 0.395, *p* = 0.056, *n* = 24). However, transfer learning was strongly influenced by motor memory retention, that is, improvements over the retention period from Last Training Trials to Posttest were associated with improvements from Last Training Trials to Transfer (*r* = 0.84, *p* < 0.001, *n* = 48), an association that was weaker for the Random (*r* = 0.48, *p* = 0.024, *n* = 24) than for the Blocked groups (*r* = 0.88, *p* < 0.001, *n* = 24; *p* = 0.007, *z* = 2.72, for the difference between correlation coefficients after Fisher r-to-z transformation). This suggests that, in general, an enhanced consolidation from Training to Posttest is strongly connected to an enhanced Training to Transfer consolidation but transfer learning was less hampered by motor memory consolidation after random than after blocked training.

Motor error quantified by EA is affected by both, predictive feedforward and responsive motor feedback. As the feedback responses typically start to compensate for feedforward errors already at 100 ms (18) and the average trial duration across groups was about 550 ms, EA should mostly reflect the feedback responses. Thus, we tested if the observed influences of training conditions also underlie feedforward motor prediction as measured by FFCF. Neither training nor sleep condition influenced motor prediction changes from Training to Posttest (retention*sleep, *F*(1,44) = 1.23, *p* = 0.274, *pEta*^*2*^ = 0.027; retention*training, *F*(1,44) = 0.56, *p* = 0.459, *pEta*^*2*^ = 0.013) or Training to Transfer (retention*sleep, *F*(1,44) = 1.37, *p* = 0.249, *pEta*^*2*^ = 0.030; retention*training, *F*(1,44) = 0.41, *p* = 0.524, *pEta*^*2*^ = 0.009). This suggests that the observed effect here is more affected by late feedback than early feedforward responses.

### Task-EEG

Explorative analysis using cluster-based statistics for a possible retention*sleep effect (with retention: Last Training Trials, Posttest; Last Training Trials, Transfer) revealed that cortical activity in all frequency bands were unaffected by Sleep vs. Wake. Thus, we focused on further task-EEG analysis regarding the training conditions (Blocked, Random).

Analysis of a possible training condition effect was restricted to the alpha band power (Fig. 3) over parietal areas according to previous work showing a linkage between training effect and force field adaptation only in alpha frequencies (19). Based on these previous findings, we defined a left-and right-hemispheric region of interest (ROIl: CP5, CP1, Pz, P3; ROIr: CP6, CP2, Pz, P4) and found a higher alpha band power for the Random compared to the Blocked groups in the Posttest and a similar effect which did not reach significance in the Transfer test, both during movement execution (Posttest, *t*(46) = -2.22, *p* =0.031, *d* = 0.642, for *t*-test of ROIl; Transfer, *t*(44) = -1.85, *p* = 0.072, *d* = 0.543, for *t*-test of ROIr). Increased alpha band values over ROIl from Training to Posttest during movement execution were associated with better task retention success (quantified by a small Training-to-Posttest difference of the motor error; Fig. 4) for participants of the Random but not of the Blocked groups (Random, *ρ* = -0.50, *p* = 0.031; Blocked, *ρ* = 0.04, *p* = 1.0, for Spearman correlations with *ρ* representing Spearmans rho). This suggests that random training of force fields affects the parietal alpha band activity of the Posttest.

**Figure 3:**
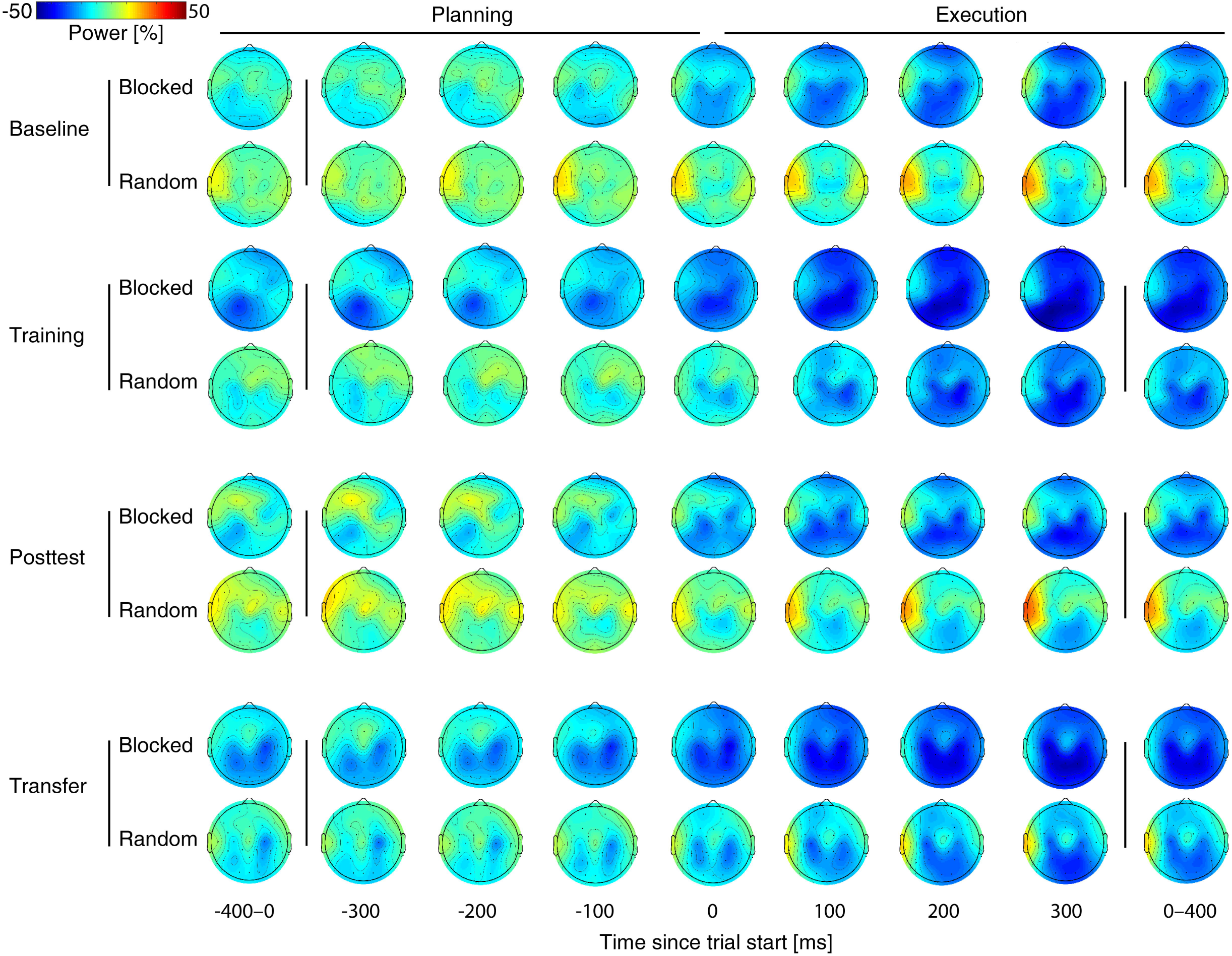
Mean alpha band power. Progress of the mean alpha band power for Blocked and Random groups during the Training session and during the Posttest and Transfer test of the Retest session. Leftmost and rightmost topographies represent the mean power across motor planning (-400–0 ms) and execution (0–400 ms) with respect to the trial start (0 ms). Other plots represent the topographical power at specific points in time (from -300–300 ms). Power values are in percentage of the average reference period 250 ms before the highlighting of a fixation cross.

**Figure 4:**
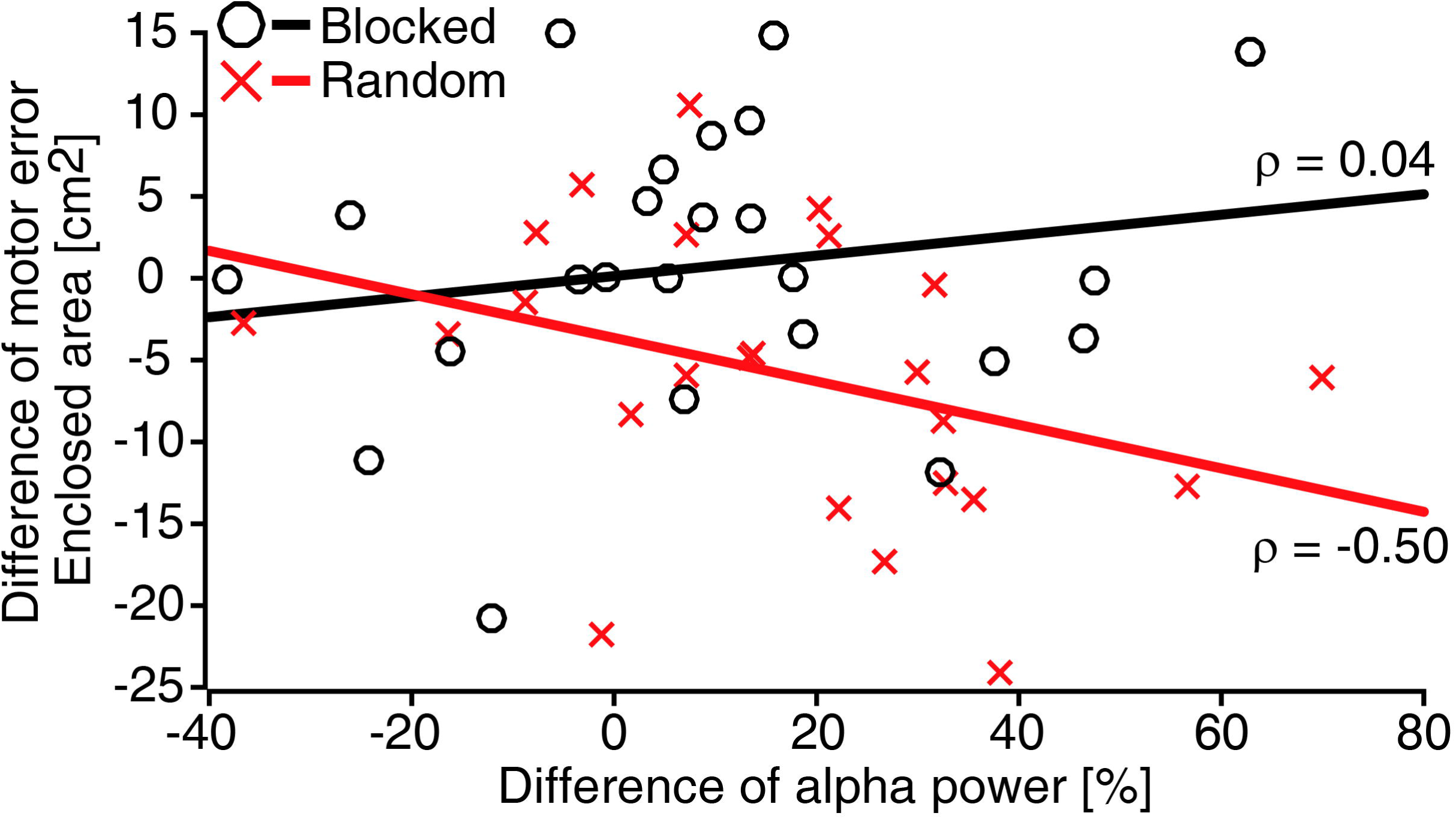
Association between Training-to-Posttest difference for motor error and alpha power. Associations between Training-to-Posttest difference of the motor error (quantified by the enclosed area) and alpha power (8–13 Hz) for ROIl (CP5, CP1, Pz, P3) during motor execution were tested using Spearman correlation. Each red cross represents the data of a single participant from the Random group and each black circle of a participant from the Blocked group. Lines represent a basic linear fit (red: Random; black: Blocked) and *ρ* represent Spearmans rho.

Furthermore, we explored if the behavioral retention effects (Training-to-Posttest, Training-to-Transfer), are predictable by EEG’s alpha band power during Training. Spearman correlations indicate positive but weak associations from Training-to-Posttest for both groups (Random, Blocked), phases (planning, execution), and ROIs (ROIl, ROIr), which did not reach statistical significance (Fig. 5). However, associations of Training-to-Transfer consolidation were strong for Blocked (*ρ* in all cases between 0.39 and 0.60) but still weak for Random groups (*ρ* between 0.04 and 0.20). These positive correlations for Blocked groups were still statistically significant after FDR correction for ROIl (Fig. 5; planning, *ρ* = 0.595, *p* = 0.025; execution, *ρ* = 0.550, *p* = 0.025) but only during trial execution for ROIr (planning, *ρ* = 0.389, *p* = 1.0; execution, *ρ* = 0.486, *p* = 0.050).

**Figure 5:**
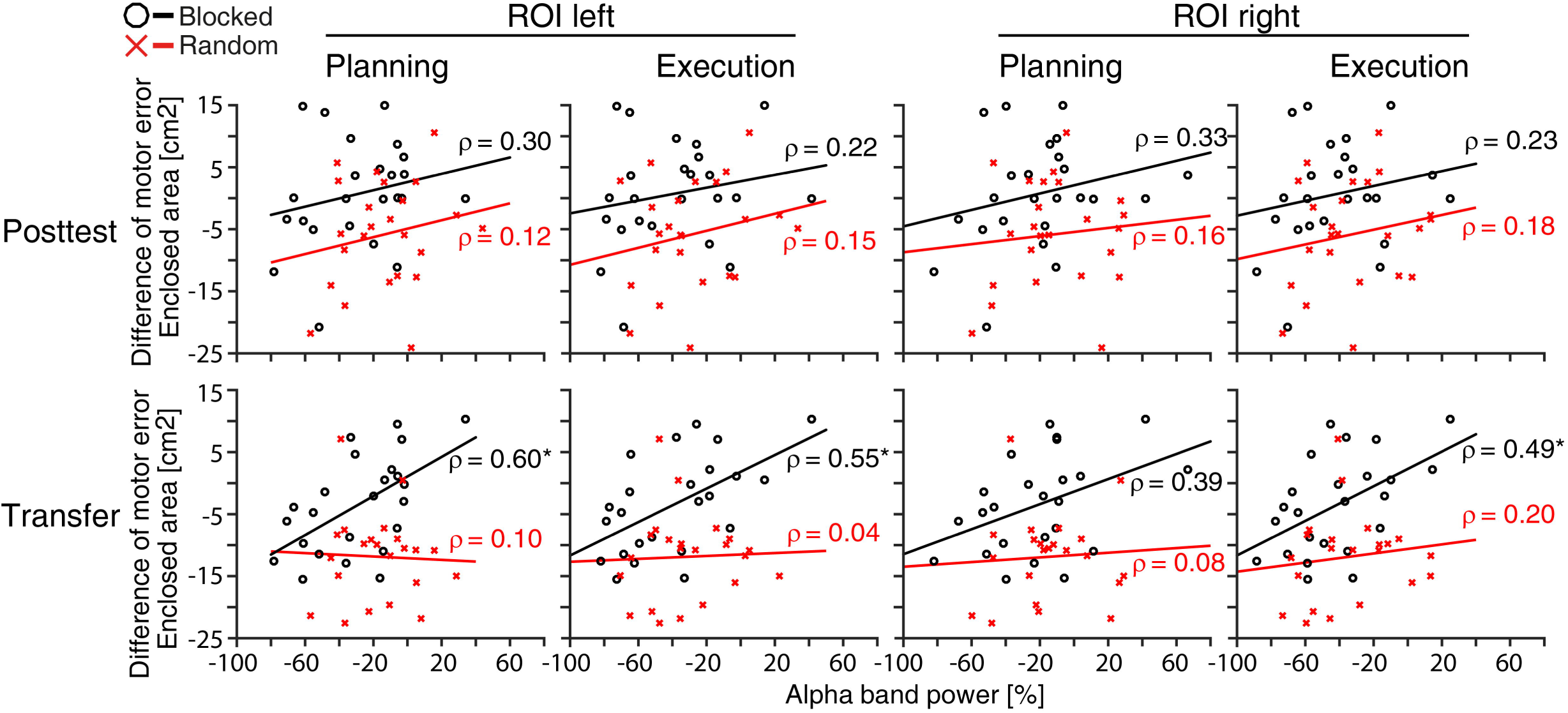
Prediction of motor memory consolidation by alpha power (8–13 Hz) during Training. The predictability of motor memory consolidation (Training-to-Posttest or Training-to-Transfer difference of motor error) is indicated. Each plot represents the data points for each participant of the Blocked (black circle) and Random (red cross) groups together with their groups’ linear fit. Asterisks indicate a significant Spearman correlation coefficient after FDR correction. Lines represent basic linear fit whereas *ρ* represents Spearmans rho.

### Sleep-EEG

Though sleep during consolidation did not improve the motor performance more than wake-time, we explored which activity during sleep could give an indication of the consolidation processes to happen during the time of sleep. None of the basic sleep stage parameters correlated with consolidation performance (Supplementary Table S1).

Consolidation success was predicted by sleep spindles and their occurrence during upstates of slow waves (Supplementary Table S2). In brief, longer sleep spindles and especially their occurrence (count and density) during upstates of slow waves were associated with improvements from Training to Posttest and Training to Transfer. This effect was most pronounced in the left parietal area (i.e., P3, corresponding to ROIl) and also indicated by more slow-wave activity (power density) during sleep-spindle enriched Stage 2 sleep. A less steep decline in oscillation frequency (i.e. less ‘chirp’ towards lower oscillation frequency) during such spindles predicted worsening from Training to Transfer, and a higher sleep spindle oscillation frequency in fronto-central areas was predictive for the deterioration of the motor performance from Training to Posttest or Training to Transfer. Higher dispersion of sleep spindles locked to the slow wave down state (in Cz), which suggests generally impaired mechanisms in the timing of sleep spindles and slow waves, predicted less improvements during Transfer compared to Training. No other sleep measures reached significant correlation with consolidation measures.

## Discussion

Our results show that Random and Blocked groups adapted to the force field conditions successfully. Subsequent consolidation was influenced by training conditions but unaffected by intervening sleep. Regarding training conditions, although Training outcome was worse for the Random than Blocked groups, all groups showed a similar motor performance in the Posttest and Random groups showed an even more pronounced motor performance compared to the Blocked groups when tested for transfer on the untrained, non-dominant hand. This improvement expressed itself in reductions in motor error (enclosed area), which is mostly affected by responsive feedback corrections, but not in the measure of motor prediction (force field compensation factor).

Our behavioral results exclude a substantial profit from sleep on the present motor adaptation task. Task performance and measures of consolidation were independent on whether participants spent awake or asleep during the intervening time between Training and Retest. Thus, our study confirms earlier findings (5) and concurs with some force field studies (20,21) claiming that consolidation of motor adaptation towards dynamic perturbations is time but not sleep dependent. Previous work suggested that force field adaptation represents a non-hippocampal dependent, implicit memory (20,22). Accordingly, our negative findings here agree with the assumption that only hippocampal dependent motor processes benefit from sleep (1). However, research showed that hippocampal damage deteriorates the benefits of unstable training conditions (16), indicating that a motor task might become hippocampal-dependent -and thus would be expected to become sleep-dependent -when trained under highly unstable conditions (1,13). However, this view is not supported by our behavioral data also showing now sleep effects for the Random groups. The exact extent to which motor adaptation after variable training becomes hippocampal-dependent is unclear and need to be elaborated in future studies.

Despite the lack of a consolidation benefit of sleep over wake retentions, the consolidation success correlated positively with sleep spindle activity during slow-wave upstates. This is at odds with the view that consolidation of motor adaptation learning is completely independent of hippocampal processes, because the coalescence of spindle and slow-wave activity during sleep is thought to benefit consolidation of hippocampal-dependent tasks in particular (22-24). Intriguingly, we found task-consolidation-associated alpha activity over parietal brain regions which matches the strong association of sleep-mediated consolidation in the same regions. This concurs with the view that cortical regions that were engaged in learning have a strong local association with spindles and slow waves in subsequent sleep (25) and predict the extent of consolidation (26). If such associations are functionally involved in the consolidation process in our data or are merely a reflection of consolidation success of other memories, not tested by our task, is unclear.

We found variable training in the motor adaptation task was predictive of consolidation benefits. This study therefore reproduced earlier findings of the contextual interference effect (12,27) in that higher training variability led to a decreased motor performance at the end of Training, but to a performance similar to that of the blocked training groups on the Posttest and to even performance benefits on the Transfer test. As previously reported (19), this gain induced by variable training is only seen in the motor error which is mostly affected by feedback responses. By contrast, motor prediction again did not show this effect. Because subjects do not receive task specific feedback during force channel trials, FFCF is only feedforward dependent (28,29). Therefore, it is likely that the motor benefits of the Random groups were facilitated by feedback corrections during movement execution evoked by the permanent regulation of random, unexpected forces during Training.

Although the Random groups ended their training worse compared to the Blocked groups, their Posttest performance was comparable. This points to unstable training conditions to either prime for better memory consolidation or the formation of memory that is more stable. Also, the generalization to the Transfer test on the left hand was more pronounced in the Random groups. The consistency of this generalization benefit over 30 trials speaks for a stable long-term memory effect. This was confirmed by the significant association of the Training performance and benefits in memory consolidation that is lower Training performance (in terms of higher motor error) from unstable training also led to better retention performance.

During Transfer testing, the participants expected a force field on their left hand that was directed in the opposite direction than force field was during Training of the right hand which explains the initially worse Transfer performance present in all groups when compared to initial Training and Posttest performance. This suggest the generalization not to take place in an extrinsic force field transformation but rather in intrinsic, mirror symmetric coordinates, that is, perturbation was expected to come from right on the right hand and from left on the left hand. This agrees with the literature (9) but contradicts the preference of extrinsic coordinates in other studies (30,31). The divergent findings of coordinate systems in use for generalization are in line with the recent assumption that representations might occur in a mixture of coordinate systems (32,33).

Paradoxically, we do not find an even more decreased initial Transfer performance for Random groups, as would be expected by a more stable intrinsic representation in this group which gives rise to predict the force field in the opposite ‘wrong’ direction during Transfer. But the opposite was the case, i.e., the Random groups showed an enhanced Transfer performance compared to the Blocked groups. There are three possible explanations for this outcome:

(1) Generalization was worse for the Random compared to the Blocked groups. Increased motor performance of the Random group might be facilitated by a weaker generalization or consolidation of the generalized memory. However, motor performance quantified by the motor prediction showed similar Transfer performances for all groups, indicating similar generalizations between groups.

(2) Random training favored the formation of a different coordinate system (or mixture of systems, 32). The results, however, do not support such explanation as motor predictions were similar between groups. In addition, inspection of single individual data revealed cues for an extrinsic force field representation in only 4 of the 24 participants of the Random groups. This was also the case in 2 of the 24 participants of the Blocked groups.

(3) Random training led to a generally increased ability to use feedback responses. This explanation is supported by the finding that only motor error, which is sensitive to feedback corrections, but not motor prediction showed an increased memory consolidation for the Random groups. In addition, the EEG data shows that parietal but not frontal areas of the brain are involved in the contextual interference effect, with the former known to be specifically implicated in sensory integration (34). However, future research should further investigate the influence of variable training on online feedback corrections in motor behavior.

Altogether, variable training leads to benefits in consolidation of a force field adaptation task. This effect is even more prominent when retention is tested on the contralateral hand. We assume that the increased consolidation after highly variable training is facilitated by an increased ability to use online feedback corrections.

Task-EEG during task performance showed that behavioral changes across the consolidation period after Random training are accompanied with a parallel increase (from Training to Posttest) in alpha band power over parietal areas, which concurs with previous findings from our lab (19). In detail, we were able to reproduce a negative correlation between changes in alpha power over contralateral parietal areas (ROIl) and motor error during movement execution. An increased alpha band power is frequently discussed as a sign of an active inhibition of the underlying cortical region (35). Therefore, a negative correlation might indicate that, for Random groups, a more accurate and, thus, better consolidated motor performance comes in parallel with an increased inhibition of parietal areas.

The results also showed that consolidation in this force field adaptation task can be predicted by the alpha power over parietal areas during Training. Blocked but not Random groups showed significant associations between Training-to-Transfer consolidation and the alpha band power. Thus, high parietal alpha power and, thus, inhibition of parietal cortical areas during Training, might favor a weaker consolidation for the Blocked but not for the Random groups. Intriguing questions arising here are whether the greater efficacy of random training specifically results from its ability to counter the disadvantage of increased parietal alpha power during training and whether parietal alpha power is connected to online feedback corrections of the motor system.

## Methods

### Participants

Forty-eight healthy, male participants recruited from the local university campus were included in the study (age 24.27 ± 0.45 yrs.). All participants were native German speakers with normal or corrected to normal vision and were tested for right-handedness by the Edinburgh handedness inventory (36). They reported not to nap habitually or have any sleep disorders and did not take any medication at the time of the experiments. Participants followed a normal sleep–wake rhythm and reported no night shifts during the 6 weeks before the experiment. Participants were instructed to keep a regular sleep schedule, abstain from caffeine-and alcohol-containing drinks for at least 2 days before and on the days of the experiments. Experimental task and task-protocol were new to the participants. All participants provided written informed consent and the study was approved by the ethics committees of the Karlsruhe Institute of Technology and the University of Tübingen.

### Apparatus and motor adaptation task

Apparatus and task stem from a previous study (see 19, for a detailed description). Participants performed point-to-point reaching movements at a robotic manipulandum (Kinarm End-Point Lab, BKIN Technologies, Kingston, Canada; Fig. 1a). The manipulandum sampled position of the handle and forces exerted on the handle at 1000 Hz. Participants’ grasped the handle and their forearm was supported by an air-sled system which enabled low friction movements. The task goal was to move a cursor on a screen – controlled via the robot handle – into a target circle (Fig. 1b). To prevent movement anticipation, each trial started with a fixation cross and the highlight-duration of this fixation cross varied randomly between 0.8 and 1.5 s. When the fixation cross changed its shape to a target circle, subjects were allowed to start their movement (no fast reaction times were required). After reaching the target, the manipulandum actively guided subjects’ hands back to the center point and provided the beginning of the next trial. In total, six targets were arranged on a circle with a diameter of 20 cm surrounding the center target. The target order was pseudo-randomized so that in every block (containing 6 movements) every target highlighted just once. In addition, within each group the target order was different for every single subject so that the mean target direction and the mean force field magnitude across all subjects was identical of each specific trial.

The manipulandum can produce forces via the handle towards subjects’ hands. In this study, we implemented three types of trials. In null field trials, no forces were produced and subjects performed movements under unperturbed conditions. In force field trials, the motors of the manipulandum were turned on and produced a velocity-dependent curl force field in clockwise direction with three different viscosity magnitudes of 10, 15, and 20 Ns/m. In force channel trials, the manipulandum produced a virtual force channel from start to target so that the subjects were only able to move along this path directly into the target (Fig. 1c). In every single trial, visual feedback about the movement time was given to ensure similar movement times across trials and subjects (< 450 ms: too slow; > 550 ms: too fast).

Offline calculations of dependent variables on the behavioral level were performed using MATLAB R2015b (MathWorks Inc., Natick, MA, United States). For null field and force field trials, we computed the motor error by using the enclosed area (EA) between subjects’ hand path and the vector joining start and target (Fig. 1c, left). This parameter was averaged over 30 trials for the Baseline, First Training Trials, Last Training Trials, Posttest, and Transfer. To quantify motor performance in force channel trials, we calculated a force field compensation factor (FFCF; Fig. 1c, right) by means of the linear regression of the measured and the ideal perpendicular force profile (29) and averaged this across each 6 force channel trials. As subjects do not receive error-feedback in these trials, this parameter reflects mainly movement prediction and, thus, feedforward mechanisms (28). From now on, the term motor error will refer to the enclosed area and the term motor prediction will refer to the force field compensation factor.

### Design and Procedures

This study compares the effects of random (unstable) vs. blocked (stable) training on motor adaptation and consolidation processes during wake vs. sleep. In a between-groups design, participants were randomly assigned to four equal sized groups (*n* = 12) of comparable age (range 18–30 yrs; *p* > 0.45, for one-way ANOVA between groups) with altered training conditions and retention periods taken place either in the night or during the day. All participants trained with their dominant right hand the motor adaptation task. The task was either trained in a random trial sequence (Random group) or in three randomized blocks, each containing a consistent field magnitude (Blocked group). Participants trained either in the morning (9 am; Fig. 1d) and were retested in the evening (8 pm; Wake-Random, WR; Wake-Blocked, WB) or, vice versa, trained in the evening and were retested the following morning after a night of sleep (Sleep-Random, SR; Sleep-Blocked, SB). The retention period between Training and Retest sessions was about 11 hours for all groups. The Wake participants spent their awake time following their usual daily activity and Sleep participants went home after Training to sleep there with polysomnographic home recordings. Retest session contained a Posttest and Transfer test, quantifying the motor performance of participants using their right (Posttest) and left (Transfer) hand (Fig. 1d).

Before Training, participants were mounted with a task-EEG and familiarized themselves with the motor adaptation task. During Familiarization, participants performed 144 null field trials with their right hand. Before Training and Posttest, participants were tested on possible confounding effects of subjective sleepiness (Stanford Sleepiness Scale, SSS, 37), mood (Positive Affect Negative Affect Scale, PANAS, 38,39), and objective vigilance (5-min Psychomotor Vigilance Task, PVT, 40).

Then, participants performed a Baseline measurement using 30 null field trials and 6 additional force channel trials. Training contained 144 force field trials followed by 6 consecutive force channel trials. All participants trained force field trials split into three force field magnitudes (10, 15 and 20 Ns/m) with a mean force field magnitude of 15 Ns/m over all trials. Random and Blocked groups trained the magnitudes under different training schedules that manipulate the training variability of those groups: the Random group trained all trials with force field magnitude switching from trial to trial in a pseudo-random order (unstable); the Blocked groups trained three trial blocks, each containing 48 trials with consistent force field magnitude, with force field magnitude switching only between the blocks. The block order was counterbalanced across participants of each group. The Wake group participants ended the Training session with unmounting of the task-EEG and were given instructions for the daytime until arrival for the Retest session in the evening; the Sleep group participants, however, were additionally prepared for the sleep-EEG and received instruction for the overnight home-polysomnography recording until the next day. The Sleep group started the Retest session with the unmounting of the sleep-EEG.

The Retest session was the same for all participants. Thereby, all participants performed a Posttest of the task with 6 force channel trials, 30 force field trials, and 6 force channel trials. All force field trials were fixed at the mean force field magnitude of the Training (15 Ns/m). Posttest was followed by a Transfer test. Transfer test had the same protocol as Posttest and participants performed the behavioral task with the non-dominant left hand. Note that the force field direction in the Transfer test was still clockwise.

### Task-EEG

To record the EEG during task performance we used the actiCHamp system with 32 active-electrodes and used the BrainVision PyCorder V1.0.6 for data recordings (Hard-and software from Brain Products, Gilching, Germany). The task-EEG was synchronized with the manipulandum via a direct link and the data was sampled at 1000 Hz. Electrodes were mounted on subjects’ heads with a cap and 29 electrodes were used for the recording of cortical activity using the international 10-10 system (Fp1, Fp2, F7, F3, Fz, F4, F8, FC5, FC1, FC2, FC6, T7, C3, Cz, C4, T8, CP5, CP1, CP2, CP6, P7, P3, Pz, P4, P8, TP10, O1, Oz, O2). The remaining three electrodes were used to record horizontal and vertical eye movements. Electrode Cz was used as the reference and Fpz as the ground electrode. The impedances of the electrodes were kept below 10 kΩ.

Offline EEG analyses were done using MATLAB R2015b (MathWorks Inc., Natick, MA, United States) and EEGLAB 13.5.4b (41). Raw data of the task-EEG was filtered first by a FIR high-pass filter with a cut-off frequency of 0.5 Hz and then by a FIR low-pass filter with a cut-off frequency of 281.25 Hz. Line noise was removed using the cleanline plugin for EEGLAB. Channels strongly affected by artifacts were removed by visual inspection and the missing channels restored using a spherical interpolation. Electrodes were re-referenced to the average reference and channel location Cz was reconstructed and appended to the data. Then, EEG data was epoched into segments of 8.5 s ranging from 6 s before to 2.5 s after trial start. Principle component analysis (PCA) was performed to compress the data to 99.9 % of the variance and, thus, deal with the reduced rank due to interpolation. Then, infomax independent component analysis (ICA, 42) was performed on the principle components. To detect bad ICA components, the components were evaluated in the spectral, spatial and temporal domain. Components showing distinct artifacts were rejected and the data was re-transformed into the channel domain.

We calculated the percentage power in the frequency domain for subsequent statistical comparisons. For this, we used complex Morlet wavelets for the frequency decomposition. We decomposed the data into 40 frequency bins ranging from 2 to 100 Hz in logarithmic space with 5 to 19 wavelet cycles changing as a function of frequency. The decomposed data was averaged over 30 trials and squared, resulting in the average power for: Baseline, First Training Trials, Last Training Trials, Posttest, and Transfer. Then, power was normalized according to the average reference period 250 ms before the highlighting of a fixation cross and the event-related desynchronization / synchronization (ERD / ERS) was calculated (43).

Data was averaged in the frequency domain into specific frequency bands: theta (4-7 Hz), alpha (8-13 Hz), beta (14-30 Hz), and gamma (30-45 Hz). The data was also compressed in the time-domain by averaging to two time windows: movement planning (-400 ms to 0 ms) and movement execution (0 s to 400 ms), where 0 s indicates the start of the trial.

### Polysomnography and sleep-EEG analyses

Standard polysomnography was assessed using a home recording system (Somnoscreen Plus, Somnomedics, Randersacker, Germany) including electroencephalography (EEG) at locations F3, F4, Fz, C3, C4, Cz, P3, P4, Pz (International 10–20 system), electrooculography (EOG) sites around the eyes, electromyography (EMG) with electrodes placed at each musculus mentalis as well as the two electrodes at each mastoids behind the ear. Fpz served as the ground electrode and Cz as the original reference. Data was digitized at 256 Hz and down-sampled to 128 Hz to facilitate computation. Offline manual sleep scoring and automatic basic sleep-EEG analysis was performed using the open-source toolbox SpiSOP (44). Data of two participants (one from the Blocked, one from Random group) were excluded for these analyses due to technical failures (n = 22). Scoring was performed by an experienced rater according to standard criteria (45) and was blind to the participant’s conditions. Sleep-EEG analyses, apart from sleep scoring, were performed on EEG channels re-referenced to the average signals from the mastoids. Sleep-EEG parameters were detected using standard settings of SpiSOP (44) based on analyses described in (46) and briefly described in the following.

#### Power spectral analyses of sleep EEG

Power spectra were calculated separately for Stage 2, SWS, non-REM and REM sleep on consecutive artifact-free 10 s intervals of non-REM sleep, which overlapped in time by 9 s. Each interval was tapered by a single Hanning-adapted window (1 s tails follow Hanning window, the other 8 s are 1) before applying Fast Fourier Transformation that resulted in interval power spectra with a frequency resolution of 0.1 Hz. Power spectra were then averaged across all blocks (Welch’s method) and normalized by the effective noise bandwidth to obtain power spectral density estimates for the whole data. Mean power density in the following frequency bands was determined: slow-wave activity (0.5–4 Hz), theta (4–8 Hz), spindles (9–15 Hz), alpha (8–12 Hz), slow spindles (9–12 Hz) and fast spindles (12–15 Hz), and beta (15–30 Hz), and log transformed (decibel) prior to statistical testing.

#### Slow waves

For the identification of slow waves, the signal in each channel during non-REM sleep epochs was filtered between 0.5 and 3.5 Hz. Next, all intervals of time with consecutive positive-to-negative zero crossings were marked as putative slow waves if their durations corresponded to a frequency between 0.5 and 1.11 Hz (zero crossings marked beginning and end of slow oscillation), yet these were excluded in case their amplitude was >1000 mV (as these were considered artifacts) or when both negative and positive half-wave amplitudes lay between -15 and +10 mV. A slow wave was identified if its negative half-wave peak potential was lower than the mean negative half-wave peak of all putatively detected slow oscillations in the respective EEG channel, and also only if the amplitude of the positive half-wave peak was larger than the mean positive half-wave amplitude of all other putatively detected slow waves within this channel. For each participant and channel, the number of slow oscillations, their density (per min non-REM sleep), mean amplitude, and slopes (down slope, the ratio between value of the negative half-wave peak and the time to the initial zero crossing, up slope, the ratio between absolute value of the negative half-wave peak and the time to the next zero crossing) were calculated.

#### Sleep spindles

For each EEG channel, the signal during non-REM epochs was filtered in a 2-Hz frequency band centered to the visually determined corresponding power peak (12 to 15 Hz range, 13.32 ± 0.11) in the non-REM power spectrum of each participant. Then, using a sliding window with a size of 0.2 s, the root mean square was computed, and the resulting signal was smoothed in the same window with a moving average. A sleep spindle was detected when the smoothed RMS signal exceeded an individual amplitude threshold by 1.5 standard deviations of the filtered signal in this channel at least once and for 0.5 to 3 s. The threshold crossings marked the beginning and end of each spindle and quantified their duration. Sleep spindle amplitude was defined by the voltage difference between the largest trough and the largest peak. Spindles were excluded for amplitudes >200 mV. We focused the analysis on fast spindles only as slow spindles power peaks could not clearly identified in too many participants. For each participant and channel’s absolute spindle counts, spindle density (per min non-REM sleep), mean amplitude, average oscillatory frequency and duration were calculated.

#### Sleep spindles co-occurring with slow wave upstates

To explore if spindles co-occurring with slow waves possess altered properties and associations with behavior, we identified slow waves that had at least one detected sleep spindle from the lowest trough (down state) to +0.5 seconds after the next positive-to-negative zero crossing (i.e., the slow wave upstate). Only the first spindle with the shortest delay to the down state was considered. Then properties of these co-occurring sleep spindles and slow waves were determined as mentioned above. In addition, the mean delay of sleep spindles to the slow wave down state as well as the standard deviation of this delay were calculated to estimate the temporal dispersion of their co-occurrence.

For an exploratory analysis of standard and fine-tuned sleep EEG parameters and their associations with memory consolidation, power density, slow wave and sleep spindle parameters (e.g. density) were averaged per electrode. Pz, F3 and F4 was each excluded from analysis in two sleep subjects and C4 in one sleep subject since these electrodes went bad during sleep EEG recording with otherwise good sleep EEG.

### Statistical analysis

We used independent two-tailed *t*-tests and mixed model ANOVAs with the within factors time (First Training Trials, Last Training Trials), retention (Last Training Trials, Posttest; Last Training Trials, Transfer), and the between factors sleep (Sleep, Wake) and training (Random, Blocked) to test differences in the motor error (EA) and motor prediction (FFCF). Data normality was tested using the Shapiro-Wilk *W*-test, and parametric or nonparametric statistical tests were chosen accordingly. For choice of appropriate *t*-tests equal variances of groups was tested using Levene’s test.

Statistical analysis of task-EEG in terms of a possible sleep effect were done using cluster-based statistics corrected by the maximum permuted cluster values (47). Therefore, mixed model ANOVAs with factors retention (Last Training Trials, Posttest; Last Training Trials, Transfer) and sleep (Sleep, Wake) were performed for every frequency band during movement planning and execution. Clusters were computed on the channel level according to p-values of the ANOVAs and the summed *F*-value for each cluster was stored as the observed statistic. Then, permutation testing was done using 1000 iterations. For each iteration, data was shuffled across both dimensions (retention, sleep), ANOVA was computed, and the maximum cluster value was stored. P-values were defined as the account of maximum permutation clusters exceeding the observed statistic divided by the number of iterations.

Furthermore, we tried to reproduce previous findings from our lab (19), targeting the neural basis for the benefits of variable training. Accordingly, independent *t*-tests between training groups (Random, Blocked) were performed to test alpha band power differences during Posttest and Transfer. In addition, motor error differences between Posttest and Last Training Trials (Training-to-Posttest) and Transfer and Last Training Trials (Training-to-Transfer) were computed for each subject on the behavioral level and correlated with the EEG data during Training using Spearman correlations.

Likewise, to find potential correlates of consolidation success within sleep parameters, we performed an explorative analysis across both Sleep groups using Spearman correlations between behavioral changes over the retention period and all sleep parameters (i.e., total sleep time [TST]; sleep onset delay; duration and percentage of TST sleep in stages like Wake after sleep onset, Stage 1, Stage 2, SWS, non-REM [i.e. SWS+Stage 2]; Power density of each sleep stage in the prominent frequency bands; parameters of slow waves, sleep spindles and their co-occurrence during the slow-wave upstates). Due to the explorative nature in the absence of a behavioral sleep effect we did not corrected these correlations for multiple comparisons.

Data and statistical analyses were performed using Matlab R2015b (Mathworks, Natick, USA) for Windows, JASP 0.8.0.1 (www.jasp-stats.org), and [R] (Windows 64bit version, 3.3.1, R Development Core Team) [The R Foundation for Statistical Computing, (www.r-project.org/foundation) 2007]. Threshold for statistical significance was set to *p* = 0.05. Multiple comparisons were either corrected by the maximum statistic (permutation test) or by the False Discovery Rate (FDR, 48). In the case of FDR, *p*-values in this study represent the FDR corrected *p*-value (49).

### Data availability

All data needed to evaluate the conclusions in the paper are present in the paper and/or the Supplementary Materials, including a Supplemental Data file (csv format) of all individual data points. Additional data related to this paper may be requested from the authors. Computer codes used to generate the results can be provided upon request. Code for sleep analyses and standard parameters is publicly available at www.spisop.org.

## Acknowledgements

We like to thank Marco Käppler for his assistance in the data recordings. This work was supported by the Graduate Funding from the German States, Deutsche Forschungsgemeinschaft (SFB 654, “Plasticity and Sleep”) and Open Access Publishing Fund of Karlsruhe Institute of Technology.

## Author contributions

B.T. and F.D.W. conceived and designed the experiment, conducted them, analyzed the results and co-wrote the paper under contribution and supervision of J.B. and T.S. All authors reviewed the final manuscript.

## Additional Information

**Supplementary information** accompanies this paper

**Competing financial interests:** the authors declare no competing financial interests.

